# Septins Coordinate with Microtubules and Actin to Initiate Cell Morphogenesis

**DOI:** 10.1101/594515

**Authors:** Diana Bogorodskaya, Lee A. Ligon

## Abstract

Many organs are formed by a process of branching morphogenesis, which begins with the formation of cytoplasmic extensions from the basal surface of polarized cells in an epithelial sheet. To study this process, we used a system of polarized epithelial spheroids, which emit cytoplasmic extensions in response to treatment with hepatocyte growth factor. We found that these extensions contain both actin and microtubules, but also septins, which are localized to microtubule bundles and appear to be important in maintaining microtubule organization. We found that these extensions are highly dynamic and form at a non-linear rate. We also demonstrated that the coordinated activity of microtubules, actin, and septins is necessary for the formation and dynamic behavior of extensions. Each cytoskeletal system plays a district role in this process, with microtubules enabling persistent growth of the extensions, actin enabling extension dynamics, and septins organizing microtubules in the extensions and supporting the extension formation. Together, our data offer insights into the dynamics of early morphogenic extensions and the distinct, but coordinated, roles of cytoskeleton in early morphogenesis.

## Introduction

Cells undergo dramatic changes in shape and structure as part of tissue and organ development, but the molecular mechanisms underlying these morphogenic events remain poorly understood. One common signaling pathway involved in many types of cellular morphogenesis is that initiated by hepatocyte growth factor (HGF). HGF is a multifunctional cytokine growth factor implicated in a wide variety of cellular processes, including morphogenesis, motility, proliferation, and resistance to apoptosis (Bando et al., 2011; Xiao et al., 2001, Rosário and Birchmeier, 2003; Steffan et al., 2011; Tokunou et al., 2001). The primary receptor for HGF is the c-Met receptor tyrosine kinase, which initiates multiple downstream signaling pathways, although Met-independent pathways have been reported as well (Devarajan, 2004). The HGF/c-Met pathway is essential to development, and knockouts are embryonic lethal. This pathway is essential in the development of organs such as placenta, kidney, lung, mammary gland and liver, and is critical for growth, regeneration, and maintenance of homeostasis in multiple organs as well (Morimoto et al., 2017; Nakamura, Sakai, Nakamura, & Matsumoto, 2011; Rosário & Birchmeier, 2003; Tokunou et al., 2001). HGF/c-Met-mediated cell motility, reorganization of the extracellular matrix (ECM), and resistance to apoptosis have also been implicated in multiple types of cancer, in which inappropriate activation of the c- MET pathway leads to aberrant cell proliferation and metastatic invasion (Nakamura et al., 2011; Scagliotti, Novello, & von Pawel, 2013).

HGF has been utilized *in vivo* to stimulate morphogenesis in a model of branching morphogenesis, a developmental pattern that leads to the formation of epithelial tubes, which are integral parts of organs such as lungs and kidney (Kim & Nelson, 2012). Several model systems have been used to dissect this process (Rosário & Birchmeier, 2003). One of the most well established models of *in vitro* tubulation involves three-dimensional cultures of Madin-Darby canine kidney (MDCK) cells. In this model, MDCK cells are seeded in a soft 3D substrate, such as collagen I, where over time they develop into a spheroid of polarized cells with the apical domains facing the lumen and the basal domains facing the ECM. In response to stimulation with HGF, some cells in the spheroids undergo a series of transformations that ultimately lead to tube formation. Individual cells undergo partial epithelial-mesenchymal transitions (pEMT) and emit extensions from the basal surface, which then elongate and undergo cell divisions to form a chain of cells. These chains continue to grow and reorganize to form cords and eventually tubes (O’Brien et al., 2004; Pollack, Runyan, & Mostov, 1998). This model has been widely used to study the process of morphogenesis and tissue regeneration, as well as more broadly, other processes involving epithelial-mesenchymal transitions (Yu et al., 2008).

Tubular morphogenesis requires coordinated changes in cell shape and invasion into the 3D ECM. At the cellular level this involves integrated reorganization of the cytoskeleton. In addition, considerable force is necessary to extend cellular protrusions into the matrix (Bordeleau, Alcoser, & Reinhart-King, 2014). Studies on downstream effectors of HGF signaling, such as ERK kinase and MMPs, have demonstrated that changes in polarity and ECM degradation are involved as well (O’Brien et al., 2004). Previous studies have shown that both the actin and microtubule cytoskeletons are necessary for this process (Gierke & Wittmann, 2012; Zhang & Vande Woude, 2003). Drugs that disrupt the actin cytoskeleton inhibit extension formation, and expression of dominant negative Rac1 or Cdc42 prevents the formation of tubes. RhoA, on the other hand, via the ROCK pathway, appears to play a role in resisting extension formation, perhaps through increasing cortical contractility (Yu et al., 2003). An early study suggested that dynamic microtubules are not required for the formation of extensions but are necessary for the transition to chains, as this process involves cell division (Yu et al., 2003). However, a more recent study showed that HGF stimulation leads to an increase in microtubule growth rates, in particular toward the basal surface of the cell and into the newly forming cytoplasmic extensions, where they often interacted with the tip and deformed the membrane. Knockdown of the +TIP protein EB1 led to a decrease in extension growth and an increase in malformed extensions with disorganized microtubules. In addition, EB1 knockdown led to defects in vesicle transport and decreased focal adhesion formation with a reduced pull on the collagen network, suggesting that the correct organization of microtubules is necessary for maintaining mechanical resistance to intracellular contractile forces and allowing for successful growth in the 3D environment (Gierke & Wittmann, 2012). Despite these studies, questions remain about the molecular mechanisms of this initial stage of morphogenesis: the formation of cytoplasmic extensions from the basal surface. Here we specifically looked at the role of microtubule bundles in extension formation and growth.

In addition to their roles in providing tracks for intracellular transport, and establishing and maintaining subcellular organization, microtubules provide structural integrity to cells and can serve as struts to support complex structures and resist or counteract actin-based compressive forces (Ingber, Wang, & Stamenović, 2014). Although individual microtubules do not have significant mechanical strength, microtubule bundles can be very strong and have the ability to resist substantial forces within the cell (Forth & Kapoor, 2017). Many proteins interact with microtubules and can potentially crosslink or bundle them. Septins, a family of GTPases that form filaments and higher order structures in the cell, have been shown to interact both directly (Bai et al., 2013) and indirectly (Bowen, Hwang, Bai, Roy, & Spiliotis, 2011; Surka, Tsang, & Trimble, 2002) with microtubules. In addition, they preferentially localize to microtubule bundles, both naturally occurring and Taxol-induced (Dolat et al., 2014; Nagata et al., 2003; Surka et al., 2002). Septins have also been shown to modulate microtubule dynamics, both through direct microtubule interaction (Bai et al., 2013) and through association with other microtubule associated proteins (MAPs), such as HDAC6 (Ageta-Ishihara et al., 2013), MAP4, (Spiliotis, Hunt, Hu, Kinoshita, & Nelson, 2008; Surka et al., 2002) and polyglutamylases (Froidevaux-Klipfel et al., 2015).

To test whether septin-associated microtubule bundles are necessary for the formation and/or growth of cytoplasmic extensions, we developed a biomimetic 3D model to investigate the early steps of branching morphogenesis in a more physiologically relevant system. We adapted the earlier model by incorporating a step to jumpstart the formation of a basement membrane around the spheroids before embedding them in a collagen gel, to more accurately model the organization of epithelial tissue. In addition, we tuned our hydrogel conditions to mimic the mechanical properties of healthy tissue (McLane and Ligon, 2015a). MDCK spheroids grown in this system respond to HGF in a stereotypical fashion similar to those in previous studies (Pollack et al., 1998). The first step of this cellular morphogenesis is the extension of cytoplasmic protrusions from the basal domain of single cells within an hour of HGF addition. Surprisingly, we found that these extensions are initially highly dynamic, often alternating between phases of growing and shrinking. These dynamics are dependent on both the actin and microtubule cytoskeletons, as are extension formation and overall growth. Microtubule bundles are prominent features of the extensions, and septins are localized to these bundles. Septin disruption blocks this co-localization and inhibits extension formation. Septin disruption at later stages of extension formation leads to malformed extensions with unbundled and disorganized microtubules, and disruption at even later stages blocks the transition from single cell extensions to chains. Together these data suggest that the early steps of cellular morphogenesis are highly dynamic and that both the microtubule and actin cytoskeleton are necessary for this process. In addition, they suggest that septin-mediated microtubule bundles are a key element in the formation and maintenance of these cytoplasmic extensions.

## Results

### HGF leads to cytoplasmic protrusions, which contain both actin and microtubules

To elucidate the mechanisms underlying epithelial morphogenesis, we developed a physiologically relevant 3D hydrogel system in which spheroids of MDCK cells are first formed in suspension in media containing a low concentration (sub-gelation) of Matrigel®, which allows extracellular matrix (ECM) components to adsorb to the spheroid surface and jumpstart the formation of the basement membrane. These spheroids with their nascent basement membranes are then embedded in a collagen-I gel of physiological stiffness, which allows the cells to efficiently develop into polarized epithelial spheroids with an intact basement membrane (McLane and Ligon, 2016). When these spheroids are treated with Hepatocyte Growth Factor (HGF), they respond in a stereotypical fashion, similar to that previously reported, by extending protrusions from the basal surface of single cells, which then elongate and undergo cell division to form chains, which will then eventually develop into tubes (Figure 1A, B). Here we focused on the initial cytoplasmic extensions that emerge from the basal membranes of individual cells. We observed considerable morphological diversity in these extensions, with some being broad at the base (encompassing the entire basal surface of the cell) and others having more narrow bases. Many tapered from the base to the tip of the extension, while others had a more consistent diameter from base to tip. Mean width at the midpoint between extension tip and base was 6.54 ± 1.99 µm (measured in a representative population of extensions (n=23) after a 24 hr incubation in HGF), but the length was more variable (discussed below). Despite this morphological variation, all extensions displayed robust actin and microtubule networks (Figure 1C), similar to previous reports (Gierke & Wittmann, 2012; Pollack et al., 1998). The microtubule network appeared especially well-organized in the narrower, less tapered, extensions in which bundled microtubules predominate (arrowheads in Figure 1C). Both microtubules and actin filaments were localized to the initial extensions formed after only one hour of HGF treatment, and both also persist in longer-lived extensions (≥18 hours HGF treatment) as well (Figure 1C).

**Figure 1.**
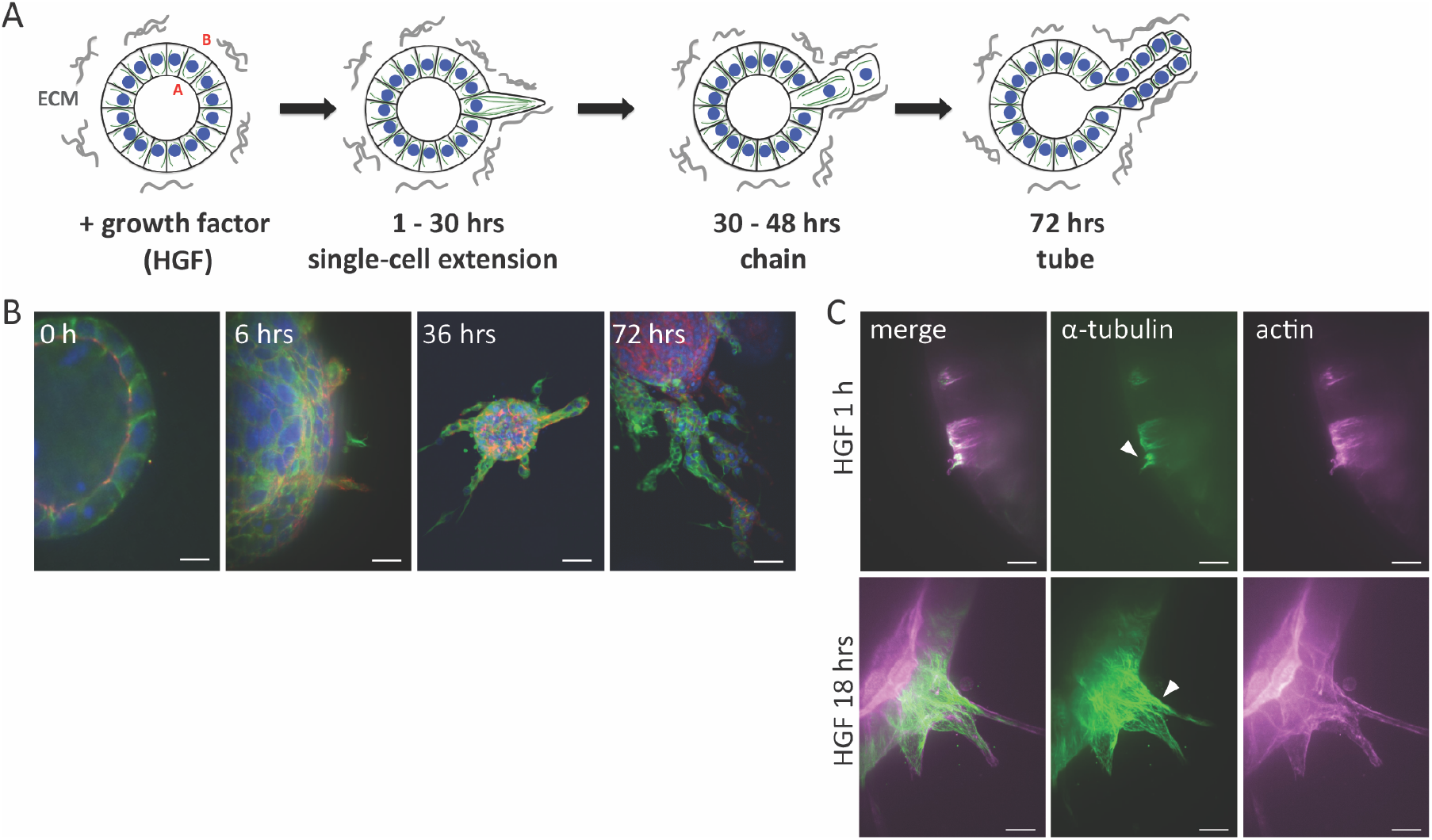
In response to HGF, MDCK cells in 3D extend cytoplasmic extensions, which contain microtubules and actin. A. Schematic of spheroids of polarized epithelial cells which undergo morphogenesis in response to exposure to hepatocyte growth factor (HGF). Letters A and B correspond to, respectively, apical and basal domains of the cells. B. MDCK spheroids exposed to HGF display cytoplasmic extensions (6 hrs), which develop into chains (36 hrs), and eventually tubes (72 hrs). α tubulin (green), zo-1 (red), F-actin (magenta), DAPI (blue). Scale = 20 µm (0 hrs), 10 µm (6 hrs), 40 µm (36 and 72 hrs). C. Extensions have both microtubules (green) and actin filaments (magenta). Scale = 20 µm. Arrowheads highlight microtubule bundles.

### HGF-induced cytoplasmic extensions are dynamic

To better understand extension growth and characterize its parameters, we used timelapse imaging of spheroids embedded in collagen gels. We imaged live spheroids using DIC microscopy. The polarized cells in the spheroids in this system begin to respond to HGF by extending cytoplasmic protrusions as early as one hour after the addition of HGF (Figure 1C), but we could not reliably detect them with DIC until ∼4 hours, so we began our recordings after 4 hours of HGF incubation and recorded for 15 hours. The rate of extension formation appears to be non-linear, with few extensions forming during the initial phase, and a significant increase in extension formation during later phases (Figure 2A, B, Supplementary video 1). The rate of extension formation was modest for the first 11 hours (phase 1), averaging approximately two visible extensions per spheroid that grow to ∼20 µm in length (although this is likely an underestimate of the total number of extensions as we will not see the extensions growing up or down in the Z-dimension). After 12-15 hours of HGF exposure, extension formation is more robust, resulting in both more extensions per spheroid and longer extensions (phase 2). This trend is accelerated after 16-19 hours of HGF exposure, resulting in even more and longer extensions in phase 3 (Figure 2B, C). In order to exclude the possibility that the higher number of extensions observed in later phases are the result of the accumulation of extensions formed earlier, we also measured extension lifetimes. We found that extensions formed in phases 1 and 2 had an average lifetime of approximately 3 hours, and this did not differ between phases 1 and 2 (Figure 2D). We could not determine the lifetimes of those extensions formed in phase 3, because our recordings ended after 19 hours. These data suggest two things: first, that the increase in the number of extensions in phases 2 and 3 is not due to the accumulation of extensions formed over the earlier phases. Secondly, they suggest that the extensions are dynamic, with relatively short lifetimes, at least during the first two phases.

**Figure 2.**
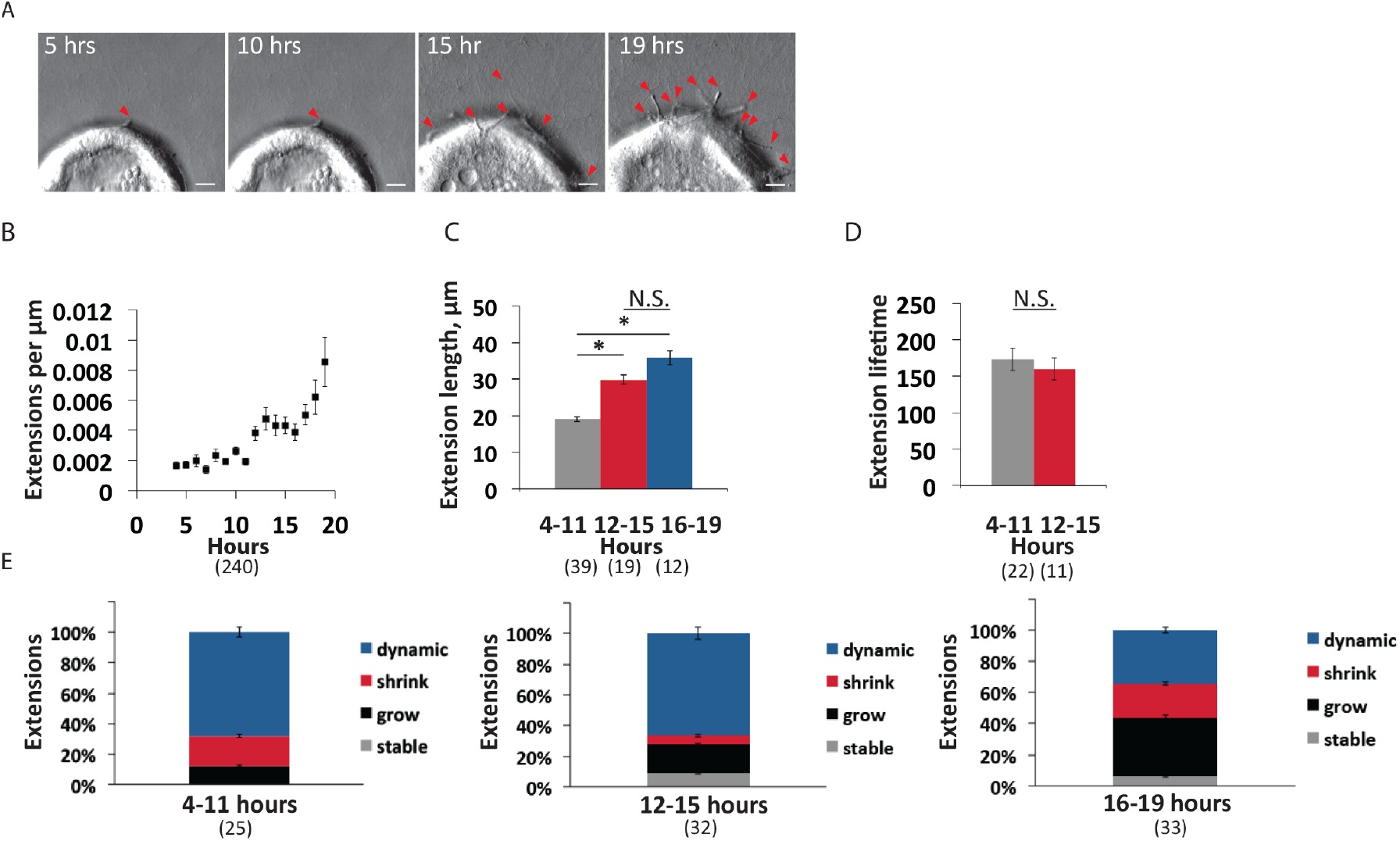
Cytoplasmic extensions are dynamic. A. Timelapse images (DIC) of spheroids (4-19 hrs after addition of HGF). Arrowheads highlight extensions. Scale = 10 µm. B. Total number of extensions per µm of spheroid circumference calculated hourly. Data represent >240 data points from 5 spheroids. Data points are mean extensions per µm of spheroid circumference at each timepoint and error bars are SEM. C. Average length of extensions during HGF treatment. Data collected from 70 extensions from 5 spheroids. Data points are means and error bars are SEM. Statistical analysis done with two-tailed t-test, p ≤ 0.05 considered significant. D. Average extension lifetime (minutes) during phase 1 (formed after 4-11 hrs HGF) and phase 2 (formed after 12-15 hrs HGF). Data collected from 33 extensions from 5 spheroids, error bars are SEM. Statistical analysis done with two-tailed t-test, p ≤ 0.05 considered significant. E. Extension dynamics during phase 1 (4-11 hrs of HGF incubation), phase 2 (12-15 hrs of HGF incubation), and phase 3 (16-19 hrs of HGF incubation). Data collected from >25 extensions from 5 spheroids. Data points are means and error bars are SEM between spheroids. Statistical analysis done using a two-tailed Fisher’s test, p≤ 0.05 considered significant.

To quantify extension dynamics, we scored each extension visible during each recording window (corresponding to phases 1, 2, and 3) and determined whether it showed persistent growth, persistent shrinkage, no change, or alternated between growth and shrinkage. In all three phases, extensions were highly dynamic. In phase 1, ∼70% of extensions switched between growing and shrinking, while smaller percentages showed persistent growth or shrinkage. None of the extensions observed in phase 1 were stable with no growth or shrinkage. The extensions seen in phase 2 were similar, although a significantly higher percentage showed persistent growth than in phase 1 (p = 0.037). By phase 3, the proportion of extensions showing persistent growth was larger still, although the proportion showing persistent shrinkage was also larger than in phase 2, resulting in a decrease in those showing both growth and shrinkage (Figure 2E).

### Extension formation and dynamics depend on both actin and microtubules

Previous studies have suggested that both actin filaments and microtubules are necessary for extension formation (Gierke & Wittmann, 2012; Pollack et al., 1998). Here we sought to determine whether an intact actin network and/or microtubule network is necessary to support the extension dynamics we observed. We incubated spheroids in HGF for 18 hours to allow for robust extension formation, and then added either an actin inhibitor (Latrunculin B) or a microtubule depolymerizing drug (Nocodazole) for two hours (Figure 3, Supplementary Videos 2-4). Both drugs resulted in a substantial disruption of the respective cytoskeletal network in extensions without obvious disruption of the other network (Figure 3A).

**Figure 3.**
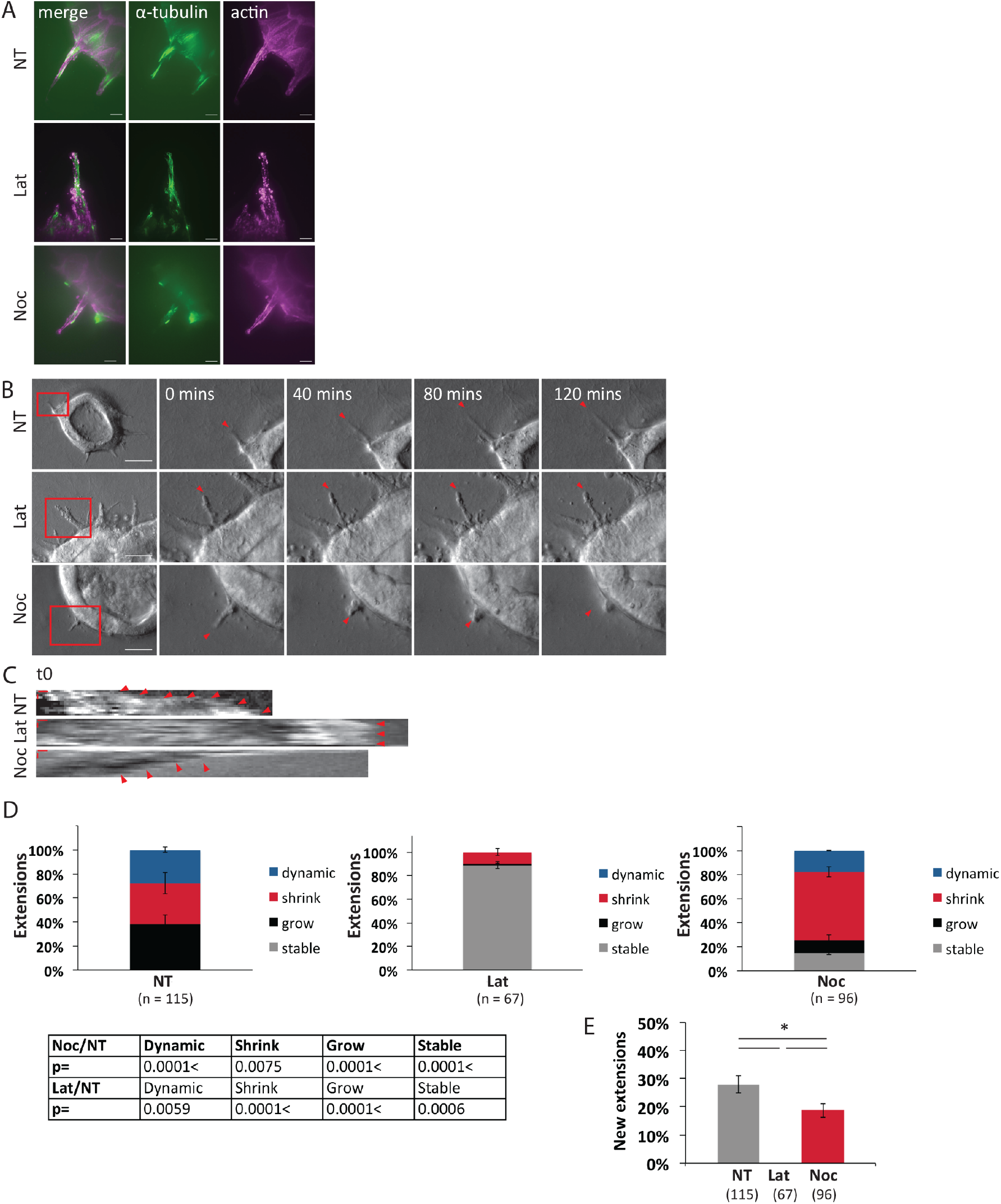
Extension growth and dynamics depend on both actin and microtubules. A. Treatment of cells with Latrunculin B or Nocodazole results in disruption of actin and microtubule cytoskeleton, respectively. Spheroids (18 hrs after addition of HGF) were treated with 1 µM Latrunculin B, 10 µM Nocodazole, or vehicle control for 2 hrs, the labeled for microtubules (anti-alpha-tubulin, green) or actin (phalloidin, magenta). B. Timelapse images (DIC) of spheroids (18 hrs after addition of HGF), with addition of 1 µM Latrunculin B, 10 µM Nocodazole, or vehicle control. Scale = 20 µm. Arrowheads highlight tips of extensions. C. Kymographs of extensions after incubation with HGF for 18 hrs, and addition of either 1 µM Latrunculin B, 10 µM Nocodazole, or vehicle control at t = 0. Horizontal scale = 1 µm, vertical scale = 20 mins. Arrowheads indicate growing extension in control (NT), stalled extension in Latrunculin B (Lat), and shrinking extension in Nocodazole (Noc). D. Extension dynamics (after incubation with HGF for 18 hrs) during 2 hr recording window after addition of either 1 µM Latrunculin B, 10 µM Nocodazole, or vehicle control. Data collected from > 67 extensions per condition, >10 spheroids per condition, in three independent experiments. Data points are means of proportions of extensions showing the specified behavior and error bars are SEM. Statistical analysis done using a two-tailed Fisher’s test, p values ≤ 0.05 considered significant. E. Percent of all extensions observed in 2 hr recording window that are newly emerged. Data collected from > 67 extensions per condition, > 10 spheroids per condition, in three independent experiments. Data points are means of shares of newly emerged extensions and error bars are SEM. Statistical analysis done using a two-tailed Fisher’s test, p values ≤ 0.05 considered significant.

We then imaged the spheroids as described previously (Figure 2), beginning approximately 20 minutes after the addition of drugs. Latrunculin treatment resulted in a dramatic increase in non-dynamic extensions, with 93% stalled. The remaining 7% showed shrinking and none grew. The extensions also showed a widening of the extension body and membrane blebbing (Figure 3B). Nocodozole treatment, on the other hand, led to a large increase in shrinking extensions and a five-fold decrease in growing extensions (Figure 3) and many showed a collapsed morphology (Figure 3B). We also tested whether new extension formation depended upon intact cytoskeletal networks by scoring de novo extension formation during the drug incubations. We found that Latrunculin treatment completely eliminated new extension formation, and Nocodazole treatment led to a 38% decrease in extension formation as well. Together these data suggest that both actin and microtubule networks are necessary for extension formation and dynamic behavior. The actin cytoskeleton appears to play a key role in initiating new extensions, but microtubules appear to be necessary for extension elongation and play an important role in extension initiation as well.

### Septins localize to microtubule bundles in extensions and are necessary for extension formation and growth

As our data suggested that microtubules are important in both extension initiation and extension growth and dynamics, and because microtubule bundles are prominent features in extensions, we sought to better understand the mechanisms underlying this microtubule organization by looking at potential crosslinkers. We focused on septins, a large family of filamentous GTPases that have been shown to associate with microtubules and may play a role in microtubule bundle formation (Bai et al., 2013; Bowen et al., 2011; Nagata et al., 2003; Surka et al., 2002). We first performed immunocytochemistry to determine if septins are localized to extensions. We used a cocktail of antibodies to septins 2 and 9, two ubiquitous subunits, and found that septins were robustly localized to cytoplasmic extensions in both the early phase of extension formation and in the later phases as well (Figure 4A). Furthermore, we found that septins co-localized with microtubules throughout the extensions, although the overlap was most prominent at the base, but did not significantly co-localize with actin filaments in the extensions (Figure 4A). Next, we tested whether disruption of the septin network would interfere with microtubule co-localization by using Forchlorfenuron (FCF), a drug that has been shown to inhibit septin subunit exchange and thus disrupt the dynamic network (Spiliotis et al., 2008). We added FCF for 4 hours to spheroids that had already been incubated with HGF for 20 hours and found a significant decrease in septin/microtubule co-localization (Figure 4A). Finally, if we added FCF to the spheroids at the same time as adding HGF, we saw a dramatic reduction in the number of extensions at 24 hours, suggesting that septins play an important role in extension initiation (Figure 4B).

**Figure 4.**
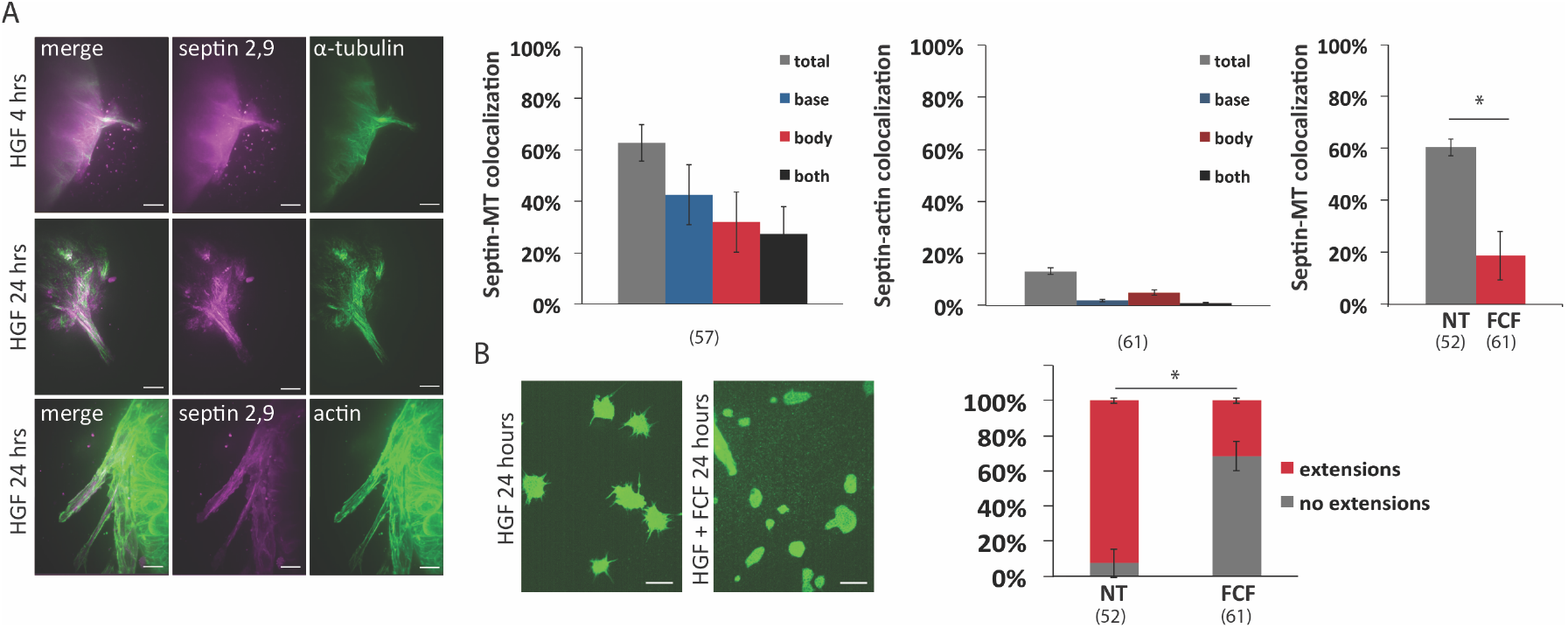
Septins colocalize with microtubules in extensions and are necessary for extension formation. A. Septins colocalize with microtubules, but not actin in extensions. Images: α-tubulin or F-actin (green), septins 2+9 (antibody cocktail) (magenta), scale = 10 µm. Septin-microtubule colocalization analysis of 24 hour of HGF dataset, from >50 extensions from >15 spheroids in 2 independent experiments. Data points are means, error bars are SEM. Scale = 10 µm. Septin disruption with 100 µM Forchlorfenuron (FCF) results in decrease in septin-microtubule colocalization. Data collected from >100 extensions in two independent experiments. Data points are means, error bars are SEM. Statistical analysis done using two-tailed Fisher’s test, p ≤ 0.05 considered significant. B. Septin disruption with 100 µM FCF inhibits extension formation. Spheroids were incubated with HGF and FCF together for 24 hrs. Cells labeled with α-tubulin (green), scale = 50 µm. Data collected from > 52 spheroids in two independent experiments. Data points are means, error bars are SEM between spheroids. Statistical analysis done using two-tailed Fisher’s test, p values ≤ 0.05 considered significant.

To further investigate the interaction between microtubules and septins in extensions, we examined the connection between septins and microtubule bundles. We found that in cells grown in 2D, septins localized to microtubule bundles, whether induced by taxol (Figure S1) or by HGF incubation (Figure 5A). We also found that when we added FCF to 3D spheroids that had been pre-incubated with HGF for 16 hours to allow extensions to form (spheroids were incubated in HGF + FCF for an additional 8 hours), resulting extensions were very disorganized in comparison to control extensions, and were often bulbous or branched (Figure 5B). The microtubules in the FCF treated extensions were often unbundled, and splayed, curly or wavy (Figure 5B), whereas in control extensions, microtubules were typically found in straight bundles.

**Figure 5.**
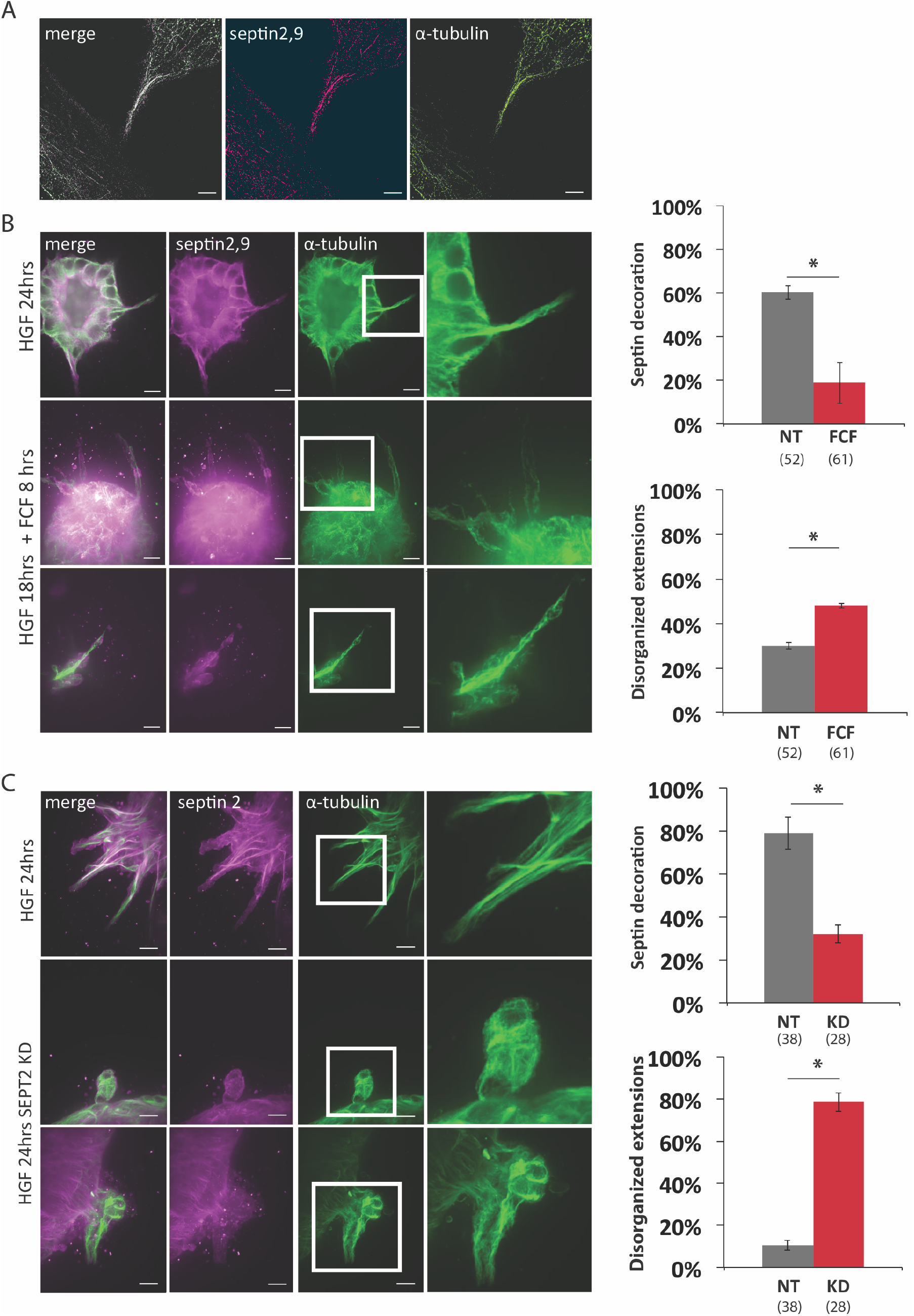
Septins localize to microtubule bundles and are necessary for the maintenance of microtubule organization in extensions. A. Super-resolution (STORM) images showing localization of septins to microtubule bundles. Images: α-tubulin (green), septins 2+9 (antibody cocktail) (magenta), scale = 5 µm. B. Disruption of septin network with FCF leads to disorganized microtubules in extensions. In extensions in control spheroids incubated with HGF for 24 hours, microtubules are predominantly straight and bundled (top row). In extensions in spheroids treated with HGF for 24 hours + FCF for the last 8 hours, microtubules are often disorganized, either unbundled and splayed (middle row), or with wavy or curly bundles (bottom row). α-tubulin (green), septins 2+9 (antibody cocktail) (magenta), scale = 10 µm. Quantification of the number of disorganized extensions (defined as those with curled, wavy or splayed microtubules) in control vs FCF treated spheroids. Data collected from > 50 extensions in two independent experiments, error bars are SEM. Statistical analysis done using two-tailed t-test, p ≤ 0.05 considered significant. C. Knockdown of septin 2 results in disorganized microtubules and misshapen extensions. In control spheroids, extensions are straight and elongated, with microtubules that are predominantly straight and bundled and exhibit robust septin decoration (top row), while in KD spheroids, extensions are rounded or irregularly shaped, with unbundled and splayed microtubules, and no septin decoration (bottom row). α-tubulin (green), septins 2+9 (antibody cocktail) (magenta), scale = 10 µm. Quantification of septin decoration of microtubule bundles, and number of extensions with disorganized microtubules (defined as those with curled, wavy or splayed microtubules) in control vs SEPT2 KD spheroids. Data collected from > 28 extensions in three independent experiments, error bars are SEM. Statistical analysis done using two-tailed Fisher’s test, p values ≤ 0.05 considered significant.

We then used siRNA-mediated knockdown of septin 2 in spheroids to further confirm the role of septins in extension organization. We chose to knock down septin 2, because members of the septin 2 family are thought to be critical to the formation of many of the septin complexes (Sellin, Sandblad, Stenmark, & Gullberg, 2011). Spheroids were transfected with septin 2 siRNA duplexes or control constructs, placed in gels for 48 hours, and then stimulated with HGF for 24 hours. Whereas the extensions in control spheroids were typically long, thin, and straight, with robust, septin-decorated microtubule bundles, the extensions in KD spheroids showed a significant decrease in the localization of septins to microtubules. In addition, the microtubules in KD extensions were curled and wavy, and the overall shape of the extensions was rounded or irregular (Figure 5C). Together, these data suggest that septins are necessary to maintain the bundled organization of microtubules in extensions, which may be necessary for extension formation and growth.

### Septins are necessary for the progression of morphogenesis

To mature into cell chains and eventually tubes (see Figure 1), extensions must first become stabilized and make productive interactions with the extracellular matrix (Gierke & Wittmann, 2012) and then the cells need to undergo cell division (Yu et al., 2003). Septins are essential to cell division (Mostowy & Cossart, 2012), so we tested whether septins were necessary for the transition from extension to chain. We incubated spheroids in HGF for 18 hours, and then added FCF for an additional 12 hours. In our system, by 30 hours of HGF exposure, many of the extensions have begun the transition to chains. We found that septin disruption during this phase of morphogenesis significantly inhibited the transition from extension to chain (Figure 6). Whereas control spheroids showed a robust population of single-cell extensions and one or more chains per spheroid, septin disruption led to a twofold decrease in average extension length (Figure 6A, B), a twofold decrease in the number of extensions per µm of spheroid circumference (Figure 6C), and a near abolishment of chain formation (Figure 6D). To confirm that the decrease in the number of chains is not due to the overall decrease in the number of extensions, we also calculated the ratio of chains to extensions and found that it was significantly decreased as well (Figure 6E). Together, these data suggest that septins are necessary for the maturation of extensions, either through structural stabilization or through cell division or both.

**Figure 6.**
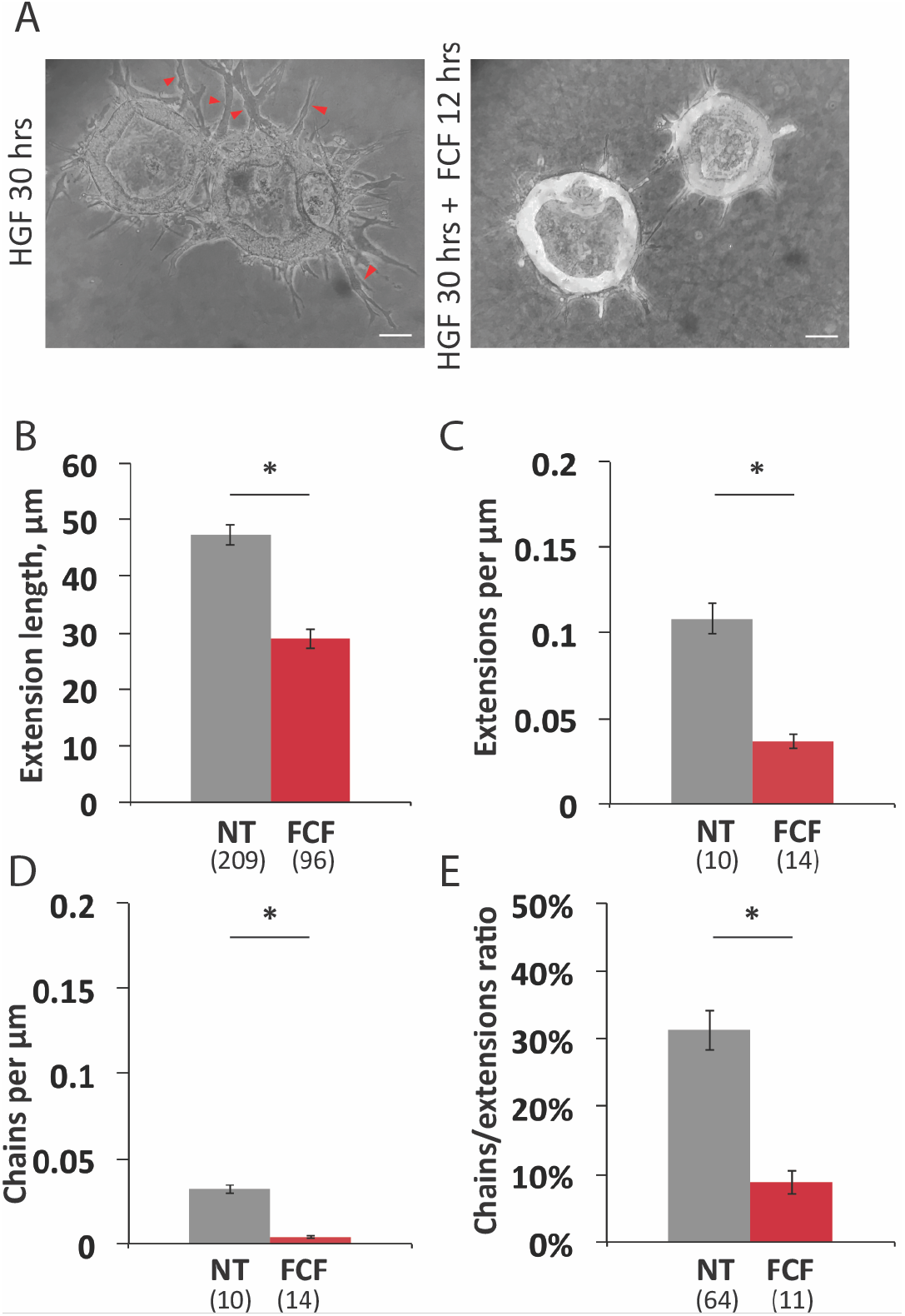
Disruption of septin network with FCF interferes with transition from single cell extensions to multicellular chains. A. Spheroids were treated with HGF for 18 hrs to allow extensions to form and then treated with HGF plus 100 µM FCF or vehicle control for an additional 12 hrs. Arrowheads indicate chains (defined as extensions containing two or more cells). Scale = 40 µm. B. Quantification of extension/chain length after 30 hrs HGF incubation (+/−FCF for the last 12 hrs). Data collected from >10 spheroids, > 90 extensions. Data points are means, error bars are SEM between spheroids. Statistical analysis done using two-tailed t-test, p ≤ 0.05 considered significant. C. Quantification of average number of extensions per µm of spheroid circumference after 30 hrs HGF incubation (+/- FCF for the last 12 hrs). Quantification of extension/chain length after 30 hrs HGF incubation (+/- FCF for the last 12 hrs). Data collected from ≥10 spheroids, ≥90 extensions. Data points are means, error bars are SEM between spheroids. Statistical analysis done using two-tailed t-test, p ≤ 0.05 considered significant. D. Quantification of average number of multicellular chains per µm of spheroid circumference after 30 hrs HGF incubation (+/- FCF for the last 12 hrs). Averages from ≥10 spheroids, ≥90 extensions, error bars represent SEM between spheroids. Statistical analysis done using two-tailed t-test, p ≤ 0.05 considered significant. E. Quantification of ratio of multicellular chains to extensions after 30 hrs HGF incubation (+/- FCF for the last 12 hrs). Averages from ≥10 spheroids, ≥90 extensions, error bars represent SEM between spheroids. Statistical analysis done using two-tailed t-test, p ≤ 0.05 considered significant.

## Discussion

Here we have used a biomimetic 3D culture of spheroids of polarized epithelial cells to elucidate the molecular mechanisms underlying cellular morphogenesis in response to HGF. We have focused on the earliest step of morphogenesis – the extension of cytoplasmic protrusions from the basal surface. We have found that these extensions are surprisingly dynamic, often alternating between bouts of growth and retraction. Consistent with previous studies, we found that both the microtubule and actin networks are necessary for extension formation and growth, but we also found that they are necessary for extension dynamics as well. In addition, we found that septin-mediated microtubule bundles are a key element in the formation and maintenance of these cytoplasmic extensions.

Intriguingly, we found that the rate of extension formation was not constant. In the first 12 hours after HGF addition, we typically observed only 1-2 extensions per spheroid. These early extensions are very dynamic, with average lifetimes of approximately three hours. The rate of extension formation increased after 12-15 hours of HGF stimulation so that there were approximately 3-4 extensions per spheroid, but these extensions were also dynamic and relatively short-lived. However, the rate of extension formation increased sharply after 15 hours of HGF exposure, and more of the extensions appeared to be persistently growing. We were not able to determine the lifetime of these extensions, but a substantial number of them continue to grow and develop into cords and eventually tubes. It is not clear what lies behind this stepwise increase in extension formation. It is possible that the accumulation of a signal, protein or post-translational modification may be necessary. New gene expression may be necessary as well. It is also possible that the acceleration of extension formation we observed might be caused by intercellular signaling rather than intracellular signaling. We noted that there appeared to be a degree of coordination between the cells of a spheroid. For example, within the same gel, some spheroids had few or no extensions, some had an average number, and some had an extremely high number (data not shown). The uniform composition of the gel and the distribution of the spheroids within the gel makes it unlikely that unequal growth factor exposure is the differentiating feature. However, it is possible that when a critical number of cells in a spheroid begin morphogenesis, other cells in the spheroid are triggered to undergo pEMT as well. Further investigation is needed into how the cells in a spheroid communicate with each other and if undergoing pEMT results in an intercellular feedback loop.

The degree of dynamicity we observed in the early extensions was also somewhat unexpected. In the early phases of morphogenesis, no extensions were static and few even showed persistent growth. Rather, most underwent periods of growth and retractions. This behavior is strikingly similar to that seen in neurogenesis, in which immature neuronal processes extend from the cell body and undergo periods of growth and retraction before one reaches a critical length and becomes the axon (Dotti, Sullivan, & Banker, 1988), suggesting that the molecular mechanisms underlying these two different types of cellular morphogenesis may be similar. Although the precise purpose of the dynamic behavior we observed in this system is unknown, it has been shown that in order to grow and stabilize, an extension must make productive adhesions to the ECM and in their absence, extensions retract (Fessenden et al., 2018; Gierke & Wittmann, 2012). Dynamic growth and retraction of cytoplasmic extensions might be a way for cells to explore the extracellular space and find favorable topology and ligands for formation of a stable adhesion. We also frequently saw sequential extensions arising from the same cell that appeared to grow along the same path through the ECM (data not shown), suggesting that newly formed extensions follow the same topological cues as the preceding ones.

Not unexpectedly, we found that microtubules and actin filaments are necessary for the formation, growth and dynamics of the extensions. The disruption of the actin cytoskeleton appears to more profoundly inhibit extension initiation than does microtubule depolymerization, probably due to its key role in remodeling the basolateral surface (Fessenden et al., 2018). However, we also observed robust recruitment of microtubules to extensions as early as we could detect them, suggesting that the cytoskeletal systems are highly temporary coordinated, and microtubules quickly grow into the nascent extensions. In addition, we saw that microtubule depolymerization did significantly inhibit extension initiation, suggesting that actin remodeling is not sufficient for extension formation. A similar pattern of microtubules populating axon-rich filopodia has been observed in neurite formation as well (Pacheco & Gallo, 2016).

Extension growth also depends on both cytoskeletal systems. Disruption of actin inhibited all dynamic activity, while depolymerization of microtubules specifically blocked persistent elongation. It is possible that actin supports all changes in extension shape, whereas microtubules, in particular microtubule bundles, provide structural support for extensions, stabilizing the shaft and enabling elongation. The precise mechanistic interaction of these two systems in this type of morphogenesis remains to be elucidated. A recent study has shown that in growing neurites, waves of actin move along the neurite shafts, transiently widening them to allow increased microtubule polymerization into the neurite, which then allows for enhanced vesicular transport to the tip to promote outgrowth (Winans, Collins, & Meyer, 2016). Perhaps a similar mechanism may play a role here.

Our initial observation of robust microtubule bundles in extensions led us to focus on microtubule crosslinkers, specifically septins. We found that septins colocalize with microtubule bundles, but not actin in the extensions, and the disruption of septins led to a decrease in colocalization. It has previously been shown that septins associate with microtubule bundles (Bowen et al., 2011; Estey, Di Ciano-Oliveira, Froese, Bejide, & Trimble, 2010; Martínez et al., 2006; Sellin, Holmfeldt, Stenmark, & Gullberg, 2011) but it remains unclear whether septins promote the formation of de novo microtubule bundles or whether they actively bundle them. Disruption of the septin network led to microtubule disorganization in the extensions. Some microtubules became unbundled, and the unbundled microtubules were typically splayed, curled, or wavy. Additionally, some of the microtubule bundles that remained were disordered, curved or wavy. This disruption affected the general shape of the extensions, resulting in rounded or irregularly shaped extensions. These finding suggest that septins organize and maintain microtubule bundles in the extensions and support extension shape. In developing neurons, septin 9 has been shown to directly bind and bundle microtubules to promote dendrite outgrowth (Bai et al., 2013), and we propose a similar mechanism may pay a role in tubulogenesis as well.

We found that the inhibition of septins abolished the formation of extensions, suggesting that they play an essential role in the morphogenic process. In addition to interacting with microtubules, septins have also been shown to interact with the actin cytoskeleton as well (Spiliotis, 2018). Although we did not observe extensive colocalization between septins and actin in the nascent extensions, it is possible that the essential role septins play may be in the remodeling of the actin cytoskeleton immediately preceding extension protrusion. A similar remodeling has been demonstrated in dorsal root ganglion neurite formation (Hu et al., 2012). On the other hand, the essential role that septins play may be due to interactions with microtubules. Septin-microtubule interactions are essential for the formation of microtubule-based protrusions that are induced by bacteria in the host cell to aid in pathogenesis (Nölke et al., 2016; Schwan & Aktories, 2016) and they have also been shown to interact with the microtubule +TIP protein EB1 to regulate microtentacle (short thin microtubule-based protrusion) formation in cancer cells (Østevold et al., 2017). Interestingly, these bacterially-induced protrusions and microtentacles show a striking morphological similarity to the cytoplasmic extensions in our system, and all are richly filled with microtubules. Finally, it is possible that septins serve to coordinate the activities of the actin and microtubule networks. It has recently been demonstrated in cortical neurons that septin disruption results in the formation of actin-rich filopodia that exclude microtubules and are unable to transition to neurites (Boubakar et al., 2017). Although the architecture of the different types of cytoplasmic extensions discussed here may vary, all show septin localization to microtubules at the base (Nölke et al., 2016; Østevold et al., 2017; Xie et al., 2007). This interaction at the base of cytoplasmic protrusions might serve to redirect existing microtubules from the cell body into the protrusions, where EB1 and other microtubule-associated proteins may direct growth along other microtubules and allow bundles to form. Alternately, septins may directly crosslink microtubules in the forming extension to aid in bundle formation. In both cases, septins facilitate the formation of microtubule bundles, which provide increased mechanical stability for these protrusions. However, more work is needed to characterize the specific roles of septins and other microtubule associated proteins involved in this process. In addition, the septin family is large with different subunits forming different complexes and a better understanding of the family in these processes is necessary.

Finally, we have shown that septins are necessary for the maturation of extensions to multicellular chains. We have hypothesized that septins play a role in extension stabilization by contributing to the formation of microtubule bundles and thereby the mechanical strength of the extension. It is possible that the failure in maturation is due to disruption of this stabilization, but it is also likely that septin disruption interferes with cell division. The transition from extension to chain requires cell division (Yu et al., 2003), and septins are known to play an essential role in cell division (Addi, Bai, & Echard, 2018), although there may be some cell-type differences (Menon et al., 2014). Further experimentation will be necessary to differentiate between these possibilities.

Our observations, combined with previous findings, lead to a more comprehensive model of the early stages of branching morphogenesis (Figure 7). Extension formation starts with a relaxation of the contractile cortical actin meshwork (Yu et al., 2003), followed by remodeling of the basal actin network (Fessenden et al., 2018). Microtubules, perhaps supported by septins, grow into the developing extensions, forming bundles that provide an extensive force, mechanical strength, and tracks for the delivery of cargoes necessary for growth (Gierke and Wittmann, 2012; our data). Nascent extensions are initially dynamic, showing phases of growth and retraction to probe the environment, and this dynamicity depends on both the actin and microtubule cytoskeleton (Figure 3). This dynamic phase is followed by stabilization, in which the extensions make a productive adhesion to the ECM, which allows for continued growth, and maturation into a chain (Fessenden et al., 2018; Gierke & Wittmann, 2012). This model suggests that the process of tubulogenesis shares a great deal of mechanistic detail with the process of neuritogenesis in developing neurons. However, further experimentation will be necessary to determine the extent of overlap in these molecular mechanisms.

**Figure 7.**
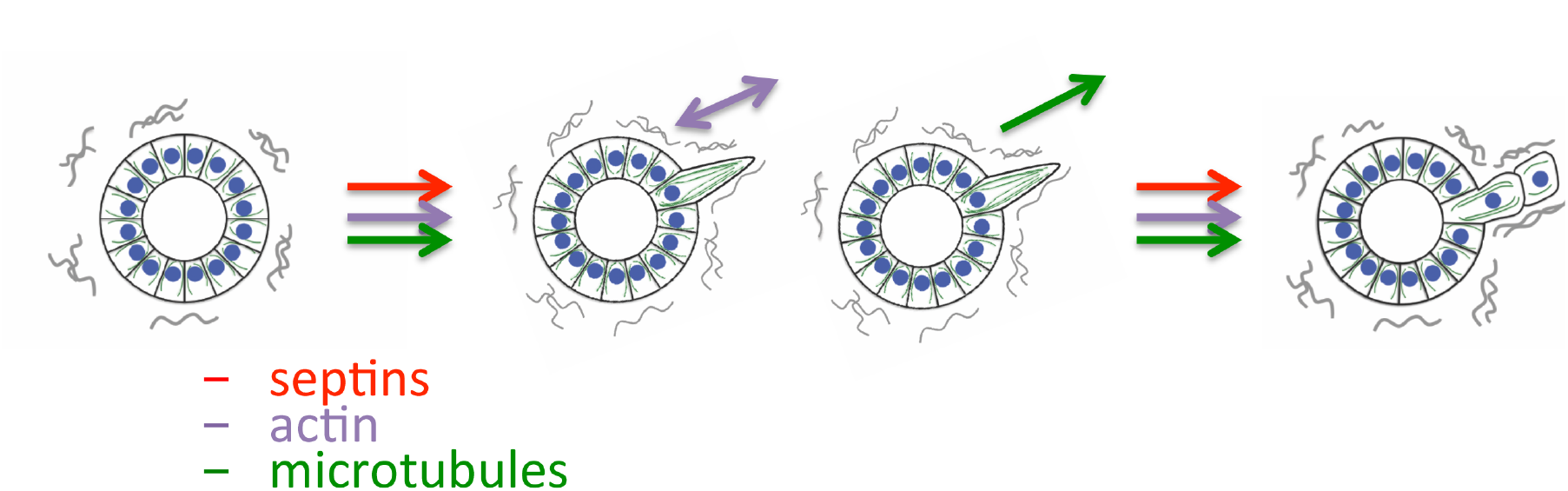
Septins, actin, and microtubules each play distinct, but coordinated, roles in early stage of cellular morphogenesis. A. Model showing coordinated involvement of septins, actin and microtubules in extension formation and dynamics. All three cytoskeletal systems are necessary for the formation of the extensions. In newly formed extensions, actin is primarily responsible for the dynamic behavior, while microtubules are necessary for the persistent growth of the extensions. And all three systems also work together for the formation of chains and continuation of morphogenesis.

## Methods

### Cells and cell maintenance

MDCK cells (product ATCC CCL34, American Type Culture Collection) were cultured according to product specifications in DMEM high glucose (Corning) supplemented with 10% fetal bovine serum (TBA), 1% penicillin/streptomycin (Corning) at 37 °C, 5% CO_2_. As cell line authentication is not readily available for canine cells, we confirmed that cells showed epithelial morphology at every step and only used cells from passages 5-25.

### Morphogenesis induction

Morphogenesis was induced by the addition of recombinant human HGF (product H1404, Sigma-Aldrich) and conditioned media from NIH/3T3 cells (product CRL-1658, ATCC). A stock solution of 3.5 µg/mL HGF in MilliQ-water was prepared and stored in 50 µL aliquots at-20°C. Aliquots were thawed on ice before use. NIH/3T3 conditioned media (CM) was prepared by growing 3T3 cells in T-75 flasks (Corning) to 70% confluence (for approximately 3 days), then collecting the culture media (3T3 cells were grown in the same media formulation as MDCK cells) and filtering it with a 0.2 µM filter. To induce morphogenesis, cells were incubated in a mixture of 25% 3T3 CM and 75% fresh MDCK media, with HGF added to a final concentration of 35ng/mL.

### Preparation of spheroids and hydrogels

Basement membrane coated spheroids of polarized epithelial cells were formed similarly to those previously described (McLane, Rivet, Gilbert, & Ligon, 2014). Briefly, 250,000 cells in 1mL of culture media were mixed with 500 µL of Matrigel® (Corning) at 3mg/mL in Opti-MEM (Corning), resulting in a final concentration of 1mg/mL Matrigel®, which is below the concentration necessary for gelation. The mixture was placed in a 15mL conical tube (Corning) which was then incubated at 37 °C, 5% CO_2_. The tubes were arranged horizontally with the lids loosely affixed to allow for air exchange. After an overnight incubation, cells were transferred to a 35mm tissue culture dish (VWR) with 1mL of culture media added. The cells were allowed to form aggregates, which then progressed to spheroids with polarized cells over 4 to 5 days. An additional 1mL of culture media was added on day 2 to supplement nutrients.

Spheroids were then embedded in a collagen-I hydrogel. To prepare the hydrogels, we adapted a method used previously (McLane & Ligon, 2015). Briefly, calf skin derived collagen I (MP Biomedicals) was mixed with neutralizing solution (0.52 M sodium bicarbonate, 0.4 M HEPES, and 0.08 N sodium hydroxide) and culture media, at the ratio of 615: 77: 308 (collagen I: neutralizing solution: media). 50 µL of the collagen-I mixture was pipetted into the bottom of the wells of an 8-chamber slides (VWR) to coat the surface and allowed to gel for 20 minutes at 37°C. A second collagen-I mixture was prepared in which the culture media was replaced with the spheroid preparation from the previous step, and 50 µL of this mixture was pipetted into the wells of the 8- chamber slide on top of the previous mixture and allowed to gel for an additional 20 minutes at 37°C. After gelation, 300 µL of culture media was pipetted into each well and the cells were maintained at 37 °C, 5% CO_2_ for up to 3 days. For some experiments, 35mm glass-bottom dishes (MatTek) were used instead of 8-chamber slides to facilitate live imaging. In this case, the volumes of the first and the second collagen-I layers were adjusted to 300 µL, and the total media volume was 2mL. For some preliminary experiments, cell aggregates were seeded in collagen I gels immediately after the overnight Matrigel® incubation step. Separate 100 µL drops of the spheroid/collagen-I mixture were placed in a 100mm dish and allowed to solidify for 45 minutes before 10mL of culture media was added to the dish. The cells were maintained at 37 °C, 5% CO_2,_ for 6–9 days, and polarized spheroids developed in 6-7 days.

### Microscopy and imaging

Unless otherwise noted, imaging was done on an inverted microscope (DMI 4000B Inverted Microscope, LEICA Microsystems) outfitted with an ORCA-ER digital camera (Hamamatsu Photonics) and a Yokogawa spinning disc confocal using Volocity imaging software (Improvision/PerkinElmer). Super resolution microscopy images were taken with an Eclipse Ti2-E microscope equipped with Nikon N-STORM 5.0 system (Nikon) using NIS-Elements software (Nikon).

Fixed cells in hydrogels (see below) were prepared for imaging by placing a gel in 35mm glass-bottom dish (MatTek), removing excess PBS with a Kimwipe (Kimtech Science), applying a drop of SlowFade® Diamond antifade mountant (ThermoFisher Scientific), allowing the gel to incorporate the antifade reagent for approximately 1 minute and then placing a glass coverslip on top to flatten the gel, optionally adding a 1g precision weight (Nasco) on top to further flatten the gel. Live imaging was done on a VivaView Fl incubator microscope with DIC optics (Olympus) using MetaMorph Software (Molecular Devices). Samples were kept at 37 °C, 5% CO_2_ and images were acquired every 10 minutes for the duration of the experiment. Low-magnification bright field images of spheroids were also acquired using a tissue culture inverted microscope (DM IL, LEICA Microsystems), with a universal smartphone adapter (Modern Photonics) and an iPhone 7 camera.

### Immunocytochemistry

Spheroids in gels were fixed with 4% paraformaldehyde in PBS for 45 min at 37°C, then permeabilized with 0.25% Triton-X in dH2O for 45 min at room temperature, washed briefly with PBS + 0.05% sodium azide (PBS-NaN_3_), and blocked for 2 hr at room temperature or overnight at 4°C in blocking solution (5% goat serum, 1% BSA, 0.05% NaN_3_ in PBS). Spheroids were then incubated with primary antibodies diluted in PBS + 0.05% sodium azide overnight at 4°C. The antibodies used were: alpha tubulin @1:500, Sigma-Aldrich product T9026; septin 2 @1:100 Proteintech product 22397-I-AP; septin 9 @1:100, Proteintech, product 10769-I-AP; ZO-1 @1:400, Life Technologies, product 33910. Antibody incubation was followed by three PBS-NaN_3_ washes, 60 minutes each. Samples were then incubated overnight with secondary antibodies (@ 1:300, Alexa Fluor, Jackson labs), and/or rhodamine phalloidin for f-actin (product P1951, Sigma-Aldrich) and counterstained with DAPI (4’,6-diamidino-2-phenylindole), followed by three 60 min washes with PBS-NaN_3_. Fixed hydrogels with spheroids were stored in 1.5mL conical tubes in PBS-NaN_3_.

Cells grown on glass coverslips (2D) were fixed with 4% paraformaldehyde in PBS for 10 min at 37°C, washed with PBS for 5 min, permeabilized with 0.05% Triton-X in dH2O for 5 min, washed with PBS for 5 min and blocked for 2 h at room temperature or overnight at 4°C in blocking solution (as above). Primary antibodies and secondary antibodies were applied in concentrations described above, with primary antibodies diluted in blocking solution and secondary antibody diluted in PBS-NaN_3_. Each antibody incubation was followed by three 5 min washes with PBS-NaN_3_. Coverslips were mounted on the glass cover slides using the ProLong® Gold antifade mountant with DAPI (Fisher Scientific).

### Inhibitors and drugs

Stock solutions of all drugs and inhibitors were diluted in DMSO (Sigma-Aldrich) and stored @-20°C. The final concentration of the drugs were the following: Latrunculin B (Cayman Chemicals) 1 µM, Nocodazole (Sigma-Aldrich) 10 µM, Forchlorfenuron (Sigma-Aldrich) 100 µM.

### Cellular morphology and colocalization assessments

Cell morphology was assessed using ImageJ (Sidhu, Beyene, & Rosenblum, 2006) or Volocity (Improvision/PerkinElmer) software packages. The number of extensions, extension dynamics, and extension length were scored or measured manually using the MTrackJ (Meijering, Dzyubachyk, & Smal, 2012) and KymoResliceWide version 0.4 (Katrukha, 2014). The organization of microtubules in extensions was assessed using an evaluation rubric: wavy extensions were classified as those in which over 50% of microtubules had bends of 20-30 degrees; curled extensions were defined as those in which over 30% of microtubules had kinks or bends of 45 degrees or more; disorganized extensions were those with curled microtubules as well as microtubules deviating more that 20 degrees from the main body of the extension (splayed microtubules). Chains were assessed manually as extensions showing the presence of two cell bodies, either in the form of visible nuclei or characteristic cell shape.

### siRNA-mediated knockdown

For siRNA knockdown, a custom IDT Oligos DsiRNA duplex for canine SEPT2 mRNA transcript variant X1 (NCBI Nucleotide accession number XM_005635975.1) was designed, with the sequence 3’ – UCACUAGCACCCAGGCACAUGCUGGCU – 3’, 3’– AGUGAUCGUGGGUCCGUGUACGACC – 5’ (Integrated DNA Technology, Coralville, IA). One-eighth of the standard spheroid preparation (250,000 cells grown over 5 days) was briefly spun down, resuspended in fresh media and spun down again. The cells were then mixed with 100uL of SE Cell Line 4D-Nucleofector® X Kit (Lonza, Basel, Switzerland) and transfected with 65nM siRNA diluted in MilliQ sterile-filtered water using the 4D-Nucleofector (Lonza, Basel, Switzerland) using the CA-152 protocol. The spheroids were transferred to a 35-mm dish of fresh warm media and left for four hours to recover post-transfection. The spheroids were then collected, briefly spun down and seeded to collagen gels as described previously. The spheroids were grown in the gel for 48 hours, then induced for morphogenesis with HGF for 24 hours and fixed.

### Statistical analysis

For statistical analysis, Microsoft Excel with the StatPlus resource pack (Release 6.1.7.0, AnalystSoft Inc.) was used. For continuous data we used a two-tailed Student’s t-test, and for categorical data we used contingency tables and Fisher’s exact test. P-values of 0.05 or lower were considered significant. For each experiment, the data were collected from at least three independent trials unless otherwise noted. Two-tailed Student’s t-test was used for continuous data analysis and two-tailed Fisher’s exact test was used to analyze the categorical data organized into contingency tables.

## Acknowledgements

The authors would like to thank Joseph Wiegartner for his contributions to the initial stages of this project, and Gerri Quinones, Brigitte Arduini and Sergey Pryshchep for technical assistance. This work was funded by National Institutes of Health grant R01GM098619.

