## Supplemental figures for "Septins Coordinate with Microtubules and Actin to Initiate Cell Morphogenesis"

**Supplementary figures**

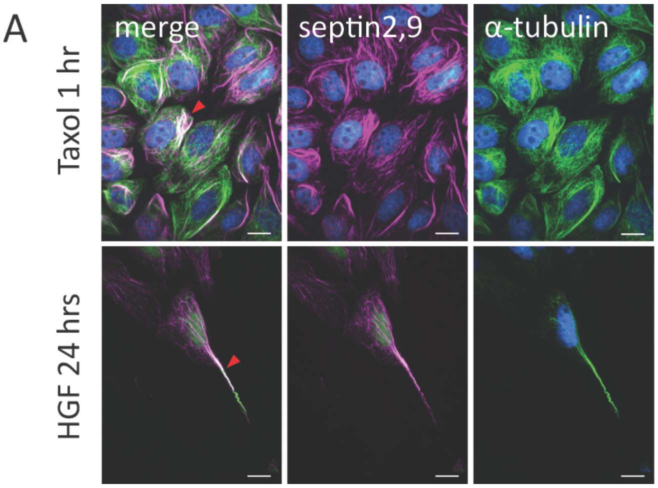


**Figure S1. Septins localize to microtubule bundles.**

A. Septins localize to microtubule bundles induced either by Taxol or by the addition of growth factor (HGF). MDCK cells grown on coverslips for 48 hours treated with either 10μM Taxol for 1 hour or with 3.5 μg/mL HGF for 24 hours, labeled for microtubules (anti-alpha-tubulin) (green) and septins 2+9 (antibody cocktail) (magenta). Arrowheads point to the areas of septin-microtubule colocalization. Scale = 10 μm.

**Supplementary videos legends**

**Video S1 Cytoplasmic extensions are dynamic and form in non-linear fashion.**
Formation of extensions Spheroid imaged for 15 hours 4 hours after the start of HGF treatment. Spheroid shows formation of multiple single-cell extensions. Length of extensions and the rate of extensions formation is increasing at later timepoints. Scale = 50 μm.

**Video S2 Cytoplasmic extension show varied dynamic behaviors.**
Spheroid imaged for 2 hours 18 hours after the start of HGF treatment. Extensions show a variety of dynamic behaviors, including growth, shrinkage, and alternating between the two, as well as formation of new extensions. Scale = 50 μm.

**Video S3 Disruption of actin network leads to loss of dynamics, abolishment of new extension formation.**
Spheroid imaged for 2 hours 18 hours after the start of HGF treatment, treated with 1μM Latrunculin B immediately prior to start of the imaging. Existing extensions are not dynamic and no new extensions are formed. Scale = 50 μm.

**Video S4 Disruption of microtubule network leads to retraction of extensions, abolishment of new extension formation.**

Spheroid imaged for 2 hours18 hours after the start of HGF treatment, treated with 10 μM Taxol immediately prior to start of the imaging. Extension appears to retract and no new extensions are formed. Scale = 50 μm.
